# BatchVaria: variance-based evaluation of batch correction with provenance tracking

**DOI:** 10.64898/2026.05.07.721996

**Authors:** Nicholas Moir, Kitty Sherwood, T. Ian Simpson

**Affiliations:** Biomedical Informatics, School of Informatics, University of Edinburgh, Edinburgh, UK; Institute of Genetics and Cancer, University of Edinburgh, Western General Hospital, Crewe Road, Edinburgh, UK

## Abstract

Assessing batch correction requires reproducible comparison of correction configurations and their effects on technical and biological variation. We present BatchVaria, an R package that integrates correction, variance profiling and reassessment within SummarizedExperiment objects. It retains original and corrected assays, records analysis provenance, variance denominators and summarisation procedures, and provides extensible interfaces for correction methods and variance engines. An application to TCGA breast cancer RNA-seq data demonstrates these capabilities across four correction configurations. Subtype variance fractions increase after every correction. Combining pooled fractions with recorded assay totals reveals that the absolute subtype contribution decreases by 4.23% under naive ComBat, whereas it is essentially unchanged (+0.06%) under subtype-protected ComBat. Under an additive plate-plus-subtype model, both `limma::removeBatchEffect` configurations have the same adjusted subtype contribution as the uncorrected assay. Their signed shared remainders nevertheless differ (−0.62% without subtype protection; +0.66% with protection), revealing a difference in attribution that the adjusted contribution alone does not capture. BatchVaria thereby supports comparison of correction configurations using absolute contributions and attribution diagnostics, rather than variance fractions alone.

**Availability:** BatchVaria is available on GitHub at https://github.com/cmpmolonc/BatchVaria.

## 1 Introduction

Batch effects arise from systematic technical differences introduced during sample processing, sequencing or data acquisition and can obscure biological signals of interest. Correction methods such as ComBat (Johnson, Li, and Rabinovic 2007, Leek *et al*. 2012) and `limma::removeBatchEffect` (Ritchie *et al*. 2015) are widely used, but their effects depend on the correction configuration and the relationship between technical and biological covariates. Datasets of molecularly heterogenous cancers such as breast can differ in molecular composition and may not reflect the population-level distribution of disease (Xie *et al*. 2020). When biological covariates are also unevenly distributed across technical batches, adjustment may remove biologically meaningful variation, making comparison of alternative correction configurations particularly important (Moir *et al*. 2025).

Tools such as variancePartition (Hoffman and Schadt 2016) and BatchQC (Manimaran *et al*. 2016) support variance modelling and batch-effect diagnostics. Reproducible comparison of correction configurations additionally requires coordinated storage of original and corrected assays, model specifications, parameter settings and evaluation results. These records must support interpretation of changes in variance attribution, rather than simply document that correction was performed.

Three details are particularly consequential. First, a biological variance fraction can increase while its absolute contribution decreases if total variance decreases proportionally more. Second, mean feature-level fractions and pooled fractions of total variance answer different questions. Third, under additive fixed-effects evaluation, adjusted Type II contributions need not sum to the full model’s explained variation; a biological contribution adjusted for technical predictors also excludes variation aligned with those predictors. Comparisons therefore require explicit variance denominators, summarisation procedures and complementary attribution diagnostics.

BatchVaria integrates these requirements within a reproducible R workflow operating directly on SummarizedExperiment objects. It combines extensible correction and variance-profiling interfaces with provenance tracking, tabular comparisons and diagnostic visualisations. We demonstrate the package using TCGA breast cancer RNA-seq data, comparing four correction configurations to illustrate how its recorded quantities support interpretation of correction outcomes.

## 2 Implementation

BatchVaria is implemented in R and is available at https://github.com/cmpmolonc/BatchVaria, alongside usage documentation and example vignettes.

### 2.1 Container and provenance

BatchVaria operates directly on SummarizedExperiment objects (Huber *et al*. 2015), maintaining compatibility with workflows accepting this Bioconductor class. Corrected assays are stored alongside their inputs, and correction and variance-profiling records are appended rather than overwritten (Fig. 1A).

**Figure 1.**
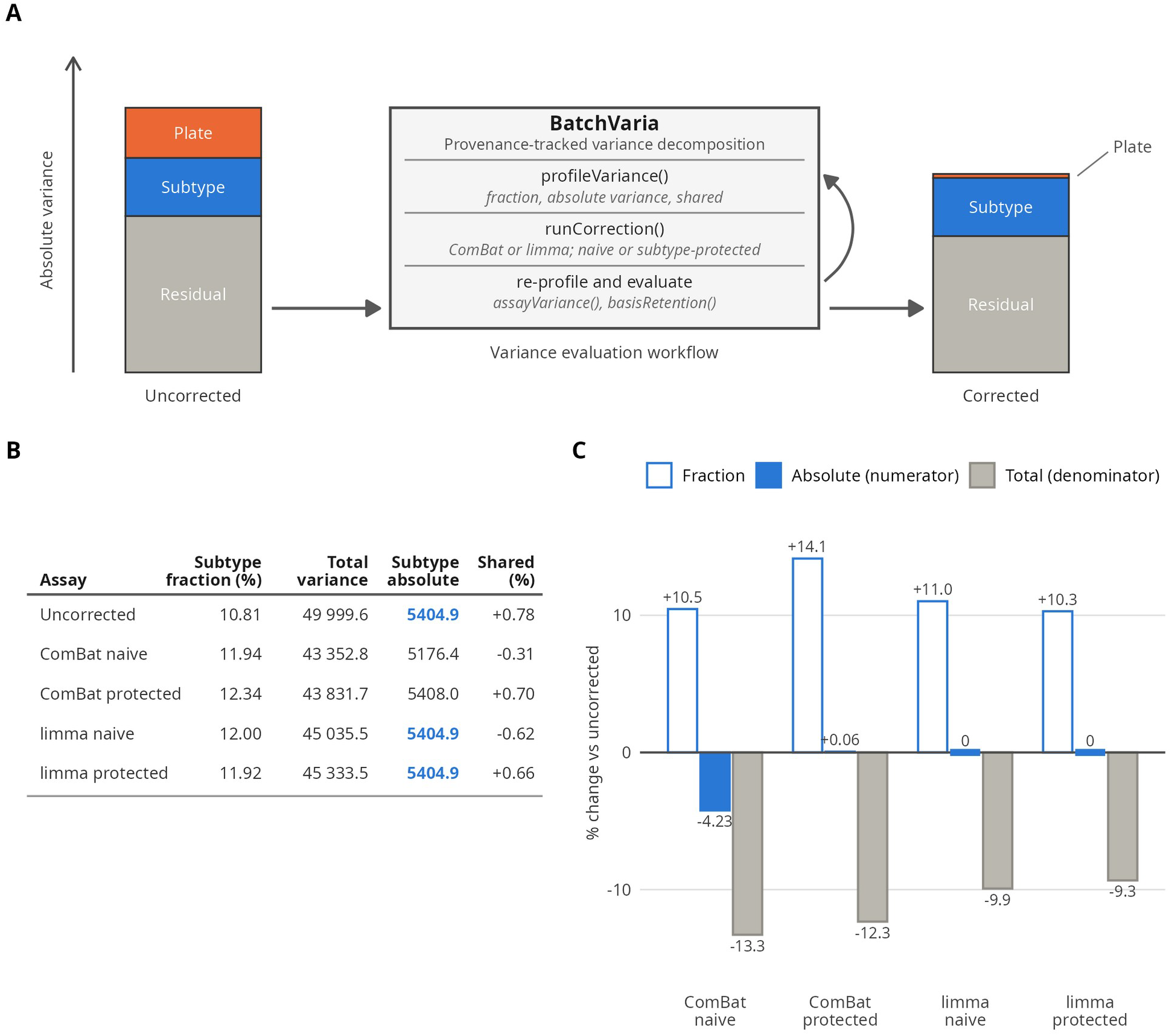
An increased subtype variance fraction can mask a decrease in its absolute contribution. Panels B and C use the anova engine with additive plate-plus-subtype models, Type II sums of squares and pooled summarisation. BatchVaria workflow and schematic (not to scale). Original and corrected assays are retained in a SummarizedExperiment, with appended correction and decomposition records, analysis provenance and assay fingerprints. The upward arrow illustrates a corrected assay undergoing re-profiling. Subtype fraction, total assay variance, absolute subtype contribution and signed shared remainder. Total variance is the sum of per-gene variances of log^2^ CPM. Bold values identify identical adjusted absolute subtype contributions in the uncorrected assay and both`removeBatchEffect` assays. Their subtype fractions differ solely because total variance changes. The adjusted subtype contribution excludes plate-aligned subtype variation; its equality does not establish equivalent preservation. The shared remainder is distinct from residual variation. Relative percentage changes from the uncorrected assay. The subtype fraction increases after all four corrections (+10.3% to +14.1%). The absolute subtype contribution decreases by 4.23% under naive ComBat, increases by 0.06% under subtype-protected ComBat, and is invariant under both`removeBatchEffect` configurations. Ticks mark zero changes. Fraction fold changes equal absolute-contribution fold changes divided by total-variance fold changes; the plotted percentage changes are therefore not additive.

Two ledgers maintain provenance. The correction ledger records the method, parameters, input and output assays, covariates specified for preservation, and total variance before and after adjustment. The variance ledger records each decomposition by assay, engine, model formula and term, with the engine configuration, including summarisation across features, retained on every result row.

Validation occurs at the point of use. Public functions check ledger structure and references to assays and sample variables. Correction methods and variance engines are checked when registered and their outputs validated before storage. Source-assay fingerprints are checked when stored results are retrieved, allowing changes to the source assay, such as subsetting or replacement, to trigger warnings when earlier results no longer describe the current data. Corrections returning an assay indistinguishable from their input are flagged. Recorded provenance is retrieved through provenance().

### 2.2 Variance profiling and summarisation

Variance profiling is performed through `profileVariance` (), using one of two engines. The anova engine fits a linear model per feature and calculates Type II sums of squares, returning term-level contributions, residual variation and a signed remainder labelled shared.

For the additive plate-plus-subtype model used here, shared is the full model’s explained sum of squares minus the sum of its adjusted term contributions. Under pooled summarisation, the remainder is expressed as a percentage of pooled total sums of squares. It is zero for orthogonal predictor blocks but can be positive or negative in non-orthogonal designs. Positive values indicate that adjusted contributions sum to less than the jointly explained variation; negative values indicate that they sum to more. The remainder diagnoses non-additive attribution and is distinct from residual variation.

Features are summarised either by averaging feature-level fractions or by pooling sums of squares. Pooled fractions can be multiplied by the corresponding assay-wide variance denominator to recover aggregate absolute contributions; unweighted mean fractions generally cannot. The selected summarisation procedure is recorded with each result.

The `variancePartition` engine wraps gene-wise mixed-model decomposition (Hoffman and Schadt 2016) and currently returns mean feature-level fractions; pooled summarisation is not implemented for this engine. A common output schema supports consistent reporting without implying equivalence between different engines or summarisation procedures.

### 2.3 Correction methods and diagnostic utilities

Corrections are applied through `runCorrection` (), which retains corrected assays and records their correction configurations. Correction methods and variance engines are extensible through `registerCorrectionMethod` () and`registerVarianceEngine` (), with built-in implementations using the same public registration interfaces as user-supplied functions. The correction methods evaluated here are ComBat and `limma::removeBatchEffect`.

`varianceTable` () reports attribution summaries and changes relative to a baseline identified from the correction ledger rather than inferred from assay names. assayVariance() returns the sum of per-feature variances, supplying the corresponding denominator for pooled ANOVA fractions.

`basisRetention` () reports the fraction of an assay’s variance lying within a reference assay’s principal-axis subspace, using all reference axes after centring and scaling each feature to unit variance in the reference. `basisProjection` () and`plotBasisProjection` () return and display coordinates in that reference basis, not principal components of the corrected assay. These diagnostics describe structural changes relative to the reference and are not measures of biological preservation.

## 3 Application to TCGA-BRCA

We used TCGA-BRCA to demonstrate a BatchVaria workflow comprising variance profiling, comparison of correction configurations and diagnostic reassessment. STAR gene counts for primary tumours (Koboldt *et al*. 2012), with PAM50 subtype annotations (Berger *et al*. 2018), were retrieved from the Genomic Data Commons using TCGAbiolinks (Colaprico *et al*. 2016). One RNA aliquot was retained per patient, with plate identifiers extracted from aliquot barcodes. Following expression filtering and TMM normalisation with edgeR (Chen *et al*. 2025), log^2^ counts per million were analysed for 30,032 features in 1,077 samples across 39 plates and five subtypes. Analyses used BatchVaria 0.99.2 with R 4.6 and Bioconductor 3.23.

Plate and subtype were associated (Cramér’s V = 0.228, P = 1.2 × 10^−4^), making their predictor blocks non-orthogonal and allowing adjusted Type II contributions to be non-additive. We evaluated ComBat and `removeBatchEffect`, each applied without subtype protection (naive) or with subtype included in the correction design. Corrected assays and their analysis records were retained alongside the original data. All assays were evaluated using the same additive plate-plus-subtype model, Type II sums of squares and pooled summarisation. Total variance was the sum of per-gene variances of log^2^ CPM. `removeBatchEffect` was evaluated as an exploratory adjustment method, not as a substitute for modelling batch in differential expression analyses.

The comparisons produced by `varianceTable` () showed that the subtype fraction increased after every correction, from 10.81% to between 11.92% and 12.34% (Fig. 1B). Combining these fractions with the total variance returned by`assayVariance` () revealed different changes in the absolute subtype contribution (Fig. 1B,C). Under naive ComBat, it decreased from 5404.9 to 5176.4, a reduction of 4.23%, while total variance decreased by 13.29%. Under subtype-protected ComBat, it was 5408.0, 0.06% above its uncorrected value. Both fractions therefore increased despite contrasting changes in absolute contribution. The naive ComBat decrease arises from scale adjustment: plate-specific mean shifts alone cannot change the adjusted subtype contribution under this evaluation model.

Both `removeBatchEffect` configurations returned the same absolute subtype contribution as the uncorrected assay, shown as 5404.9 in Fig. 1B. This equality follows because subtracting fitted effects within the plate-design span leaves the adjusted subtype contribution unchanged. The subtype fraction was nevertheless 10.81% before correction, 12.00% after naive correction and 11.92% after subtype-protected correction, with these differences arising entirely from changes in total variance. Crucially, the adjusted statistic excludes plate-aligned subtype variation and cannot detect its removal by naive correction. Equality of this statistic therefore does not establish equivalent preservation of subtype structure.

The shared term recorded by `profileVariance` () supplied complementary information. The signed shared remainder was +0.78% before correction, -0.31% and -0.62% after naive ComBat and naive `removeBatchEffect`, and +0.70% and +0.66% after their subtype-protected counterparts (Fig. 1B). For naive `removeBatchEffect`, the non-positive sign follows algebraically from removing the plate projection; ComBat has no corresponding sign constraint. These observations illustrate the diagnostic’s use in comparing configurations, not a general sign rule for identifying successful correction.

Recording the summarisation used by `profileVariance` () also allowed its effect on the reported fractions to be assessed: in the uncorrected assay, mean-of-fractions summaries were 21% lower for subtype and 39% higher for plate than the corresponding pooled summaries. `basisRetention` () reported 100% retention for both `removeBatchEffect` configurations, as expected for a correction that subtracts linear combinations of the original samples, and above 99% for both ComBat configurations, despite their differing absolute subtype contributions. These outputs demonstrate why summarisation and structural diagnostics must be interpreted alongside the quantities they describe.

Finally, dividing the pooled plate and residual fractions by their respective degrees of freedom and taking their ratio gave a plate-to-residual mean-square ratio of 3.14 before correction and below one afterwards. These values describe attenuation under the evaluation model. Because estimating and removing plate effects from the same data also removes noise aligned with plate, the post-correction ratios do not independently establish that batch effects are absent.

## 4 Discussion

BatchVaria integrates correction, evaluation and provenance tracking in a single workflow for comparing correction configurations. By retaining assays, model specifications, variance denominators and summarisation procedures together, it supports reproducible interpretation and reporting of alternative analyses. It complements existing correction methods rather than introducing a new correction algorithm or an automatic ranking criterion.

The demonstration supports three practical uses of the package: reporting total variance alongside fractions, using pooled summaries for aggregate absolute comparisons, and examining the signed shared remainder when evaluating covariate protection. The remainder complements adjusted contributions but is neither a direct measure of removed biological signal nor a sign-based success criterion. Similarly, reference-basis retention describes structural relationships rather than biological preservation.

The demonstrated application is limited to one dataset and one batch variable, and its analytical conclusions depend on the specified evaluation model. Pooled summaries also give greater weight to high-variance genes and may obscure effects concentrated in a small subset of biologically informative features. Where preservation of particular biological effects is essential, these summaries should be complemented by gene-level and downstream analyses. BatchVaria supplies the coordinated records and diagnostics needed to support those decisions without prescribing a universally optimal correction.

### Reproducibility

All code to reproduce analysis contained in this manuscript is available online: github.com/cmpmolonc/BatchVaria-AppNote.

